# SMARTFlexDB: a paired apo-holo RNA structure database and analysis platform for RNA conformational flexibility and small-molecule recognition

**DOI:** 10.64898/2026.10.11.757724

**Authors:** Shiyu Jiang, Jinhang Wei, Amirhossein Taghavi, Gustavo Seabra, Jessica L. Childs-Disney, Chenglong Li, Matthew D. Disney, Yanjun Li

## Abstract

RNA structures can undergo substantial conformational changes upon small-molecule binding, yet existing structural resources provide only static RNA-ligand snapshots without connecting ligand-bound (holo) RNA structures to corresponding ligand-free (apo) conformations. SMARTFlexDB is a curated database and analysis platform that uses paired apo-holo RNA structures to characterize conformational differences and RNA–small-molecule recognition. Candidate pairs were identified from the Protein Data Bank by matching RNA chains with at least 90% sequence identity and coverage, and fully conserved binding pocket nucleotides. The current release contains 434 binding pockets from 266 holo structures, linked to 186 apo structures to yield 1,187 pocket-level apo-holo pairs organized into 734 database entries, spanning 150 small molecules and 23 RNA functional classes. Each entry integrates RNA sequence and secondary-structure annotations, apo and holo three-dimensional structures, RNA-ligand atomic interactions, and quantitative descriptors of global and local conformational changes. An interactive web interface enables users to explore sequence alignments and structural overlays, compare secondary structures and RNA motifs, and upload paired apo-holo structures for customized analysis. By linking ligand-bound and ligand-free RNA structures, SMARTFlexDB provides a robust foundation for investigating RNA structural flexibility and supporting RNA-targeted small-molecule drug discovery. The database and analysis platform are available at https://aidd.rc.ufl.edu/app/smartflexdb/.

**GRAPHICAL ABSTRACT:** 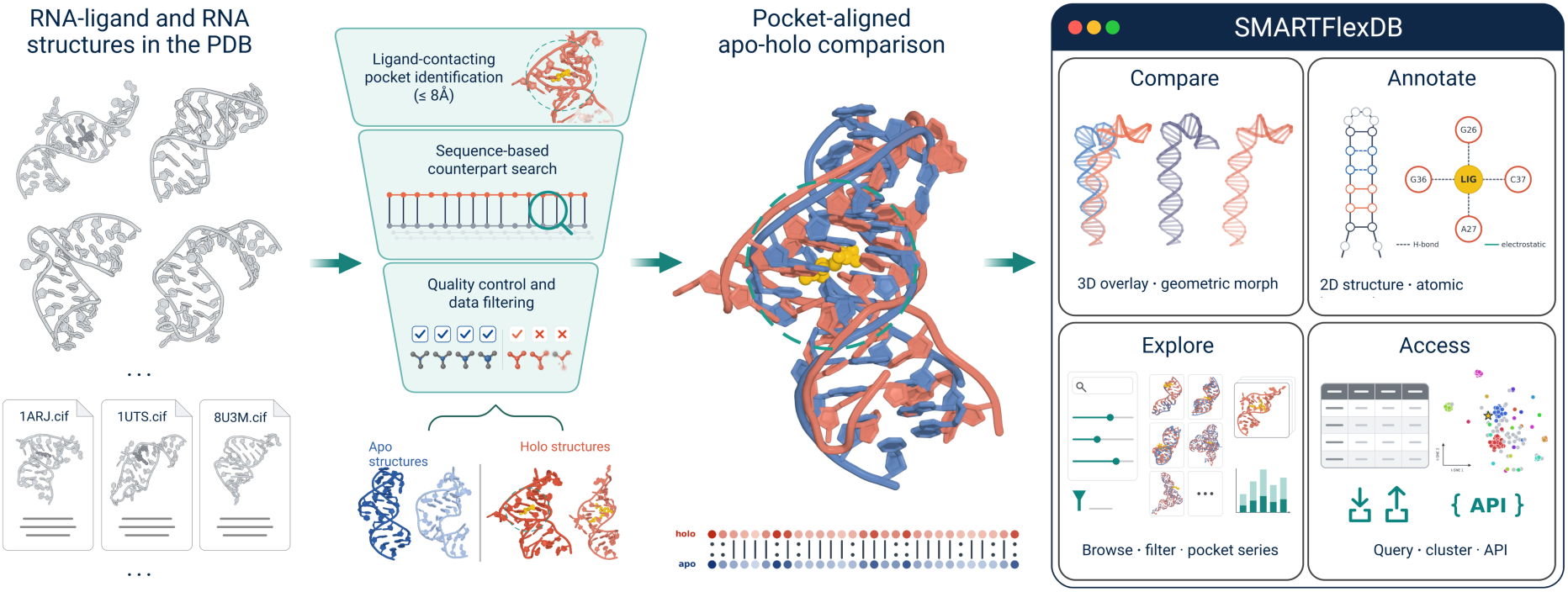

## INTRODUCTION

RNAs are important therapeutic targets, and small molecules that selectively engage functional RNA structures have emerged as a promising modality for modulating RNA activity^1–3^. However, RNAs often populate multiple conformational states that are coupled to gene regulation, catalysis, translation and molecular recognition^4,5^. Small molecules may recognize pre-organized or transient pockets and stabilize particular structural states^6–8^. Consequently, RNA-ligand recognition cannot always be understood from a single static structure. Direct comparison of experimentally determined ligand-free (apo) and ligand-bound (holo) states is required to distinguish conserved structural features from ligand-associated differences in pocket geometry, nucleotide position and base pairing^9,10^.

Paired structural information is becoming increasingly important for computational RNA drug discovery, particularly as machine learning and structure-based methods for RNA binding site identification, ligand ranking, and RNA-small molecule recognition continue to rapidly advance^11–14^. These developments demonstrate the value of three-dimensional structural information for RNA-ligand interaction prediction and virtual screening^15,16^. However, most structure-based workflows represent an RNA target using a single conformation, often a ligand-bound structure. Because RNA binding pockets can adopt different conformations in the apo and holo states^17^, methods developed or evaluated using holo structures may not perform similarly when applied to ligand-free apo structures^13,18^. Recent machine learning studies further showed that agreement between apo- and holo-state pocket-druggability predictions decreases as the structural differences between the two states increase^19,20^. Systematically paired apo-holo structures can address this limitation by enabling direct measurement of ligand-associated conformational changes and comparison of pocket properties between the two states. Such datasets can therefore be used to determine which structural features remain accessible before ligand binding and to benchmark computational methods on matched apo and holo representations of the same RNA target^11,21,22^.

RNA-focused small-molecule libraries and databases have expanded over time, providing foundational knowledge and design principles for RNA-targeted ligand discovery^23^. Representative publicly available resources include SM2miR^24^, a database of the experimentally validated small molecules that affect microRNA expression; Inforna 2.0^25^, a database of experimentally determined RNA-small molecule interactions that enables sequence-based design; NoncoRNA^26^, a database of experimentally supported non-coding RNAs and drug targets in cancer; R-BIND 2.0^27^, a database of bioactive RNA-targeting small molecules and associated RNA secondary structures; and R-SIM^28^, a database of binding affinities for RNA-small molecule interactions. However, most RNA-ligand interaction resources provide only RNA sequence and/or secondary-structure information, without incorporating the three-dimensional atomic structural context of RNA that defines ligand recognition. In contrast, HARIBOSS^29^ compiles experimentally determined three-dimensional RNA-small molecule complexes and annotates their ligand, pocket and interaction properties. Despite their complementary scopes, existing RNA structural resources do not systematically capture and analyze the conformational changes associated with small-molecule binding. In particular, ligand-bound RNA structures are not routinely linked to their corresponding ligand-free states, making it difficult to determine how the structure and properties of a binding pocket change upon ligand recognition.

Systematically pairing apo and holo structures provides a direct framework for identifying and visualizing these changes, revealing which pocket features are pre-existing in the apo state and which emerge or reorganize upon ligand binding. Such comparisons can provide mechanistic insight into RNA-ligand recognition and establish experimentally derived benchmarks for evaluating structure-based computational methods across different RNA conformational states. Furthermore, paired apo-holo analyses have been established for proteins through resources such as PLINDER^30^, which links protein-ligand complexes to corresponding unbound structures and supports the development and rigorous evaluation of cutting-edge computational methods for protein-ligand interaction modeling^31^, molecule design^32,33^ and broader protein-targeted drug discovery. However, no equivalent resource has been available for RNA, especially given that RNA molecules are highly flexible and frequently undergo conformational rearrangements upon ligand binding. Here, we present SMARTFlexDB (**<u>S</u>**mall **<u>M</u>**olecule **<u>A</u>**pproaches to **<u>R</u>**NA **<u>T</u>**argeting **<u>Flex</u>**ibility **<u>D</u>**ata**<u>B</u>**ase), a curated database of experimentally determined apo-holo RNA structural comparisons. SMARTFlexDB identifies ligand-bound RNA structures in the Protein Data Bank and searches for sequence-identical or highly similar ligand-free RNA counterparts. Candidate apo-holo pairs then undergo sequence- and pocket-level verification to confirm nucleotide correspondence, binding-site conservation, and the absence of ligand in the entire apo structure (Figure 1a). Each accepted apo-holo comparison is then characterized across four interconnected dimensions: RNA sequence and secondary structure, three-dimensional conformational changes, ligand chemical properties, and RNA-ligand interactions. Each database entry represents a unique apo-holo PDB pair, allowing each structure to be linked to multiple matched counterparts. The database also organizes complexes containing different ligands bound to a common RNA region into multi-ligand pocket groups, and provides sequence-, structure-, and pocket-based clustering to support redundancy-aware analyses. SMARTFlexDB is accessible through an interactive web interface and programmatic access, while an online query service enables users to apply the same comparative workflow to their own apo and holo structures.

**Figure 1.**
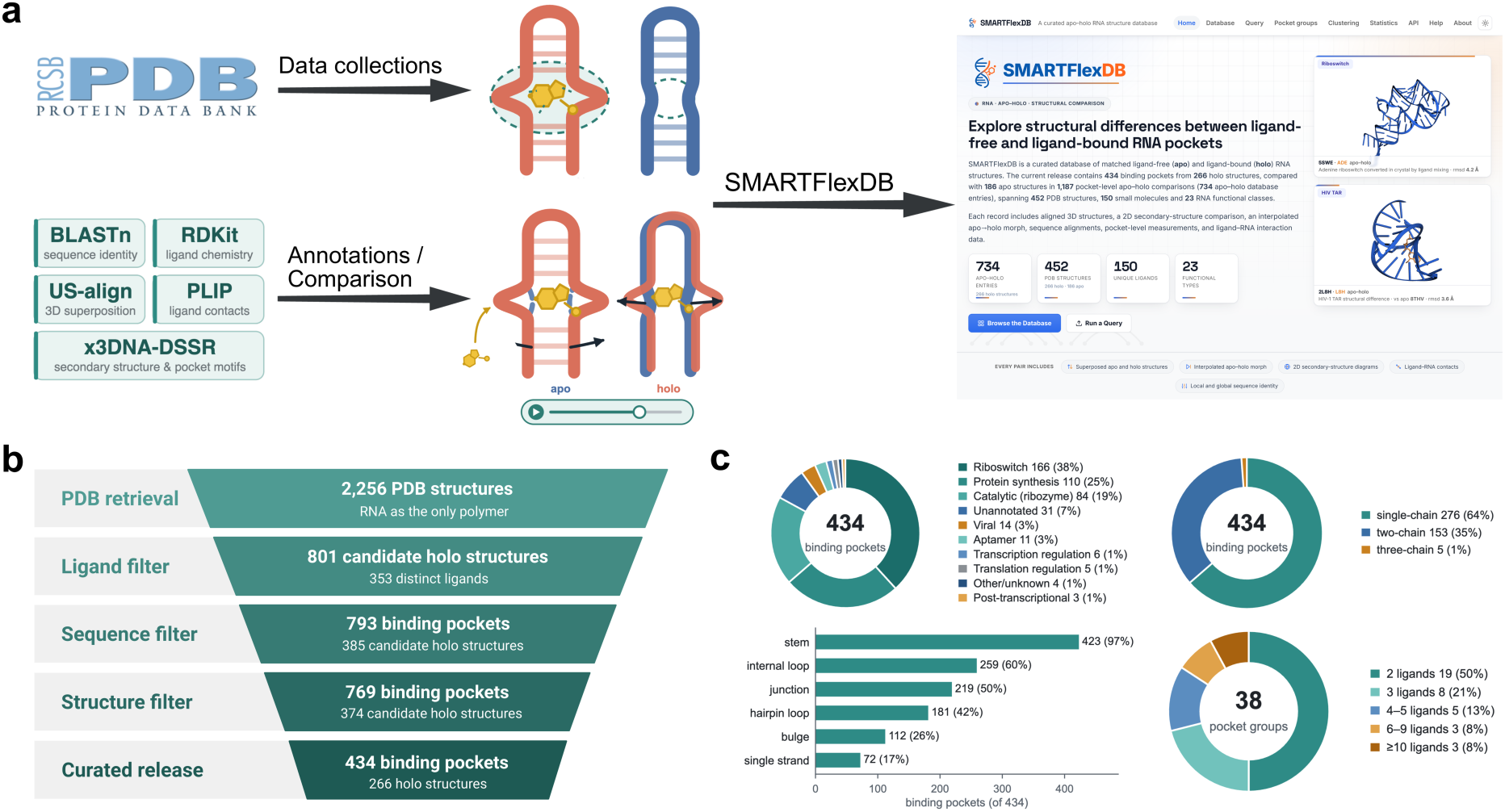
Overview of SMARTFlexDB construction and dataset composition. **(a)** Schematic overview of the SMARTFlexDB workflow. RNA structures are collected from the RCSB PDB, candidate ligand-bound and ligand-free counterparts are identified and verified, and accepted apo-holo pairs are subjected to sequence, structural, ligand and RNA-ligand interaction annotation and comparison before integration into the SMARTFlexDB web platform. **(b)** Data-curation funnel from the initial PDB retrieval through ligand, sequence and structure filters to the curated release, yielding 434 RNA binding pockets from 266 holo and 186 apo structures, forming 1,187 pocket-level comparisons organized into 734 apo-holo database entries. Intermediate counts indicate the number of PDB structures, candidate holo structures, distinct ligands and RNA binding pockets retained at each stage. **(c)** Composition of the curated dataset across coarse-grained RNA functional categories, number of RNA chains contributing to each binding pocket, prevalence of annotated RNA secondary-structure elements, and multi-ligand pocket group size according to the number of distinct ligands represented within each multi-ligand pocket group. The figure was created in BioRender. Jiang, S. (2026) https://BioRender.com/t8y5xht.

## MATERIAL AND METHODS

### Data collection

SMARTFlexDB compiles experimentally determined ligand-bound (holo) and ligand-free (apo) RNA structures retrieved from the RCSB Protein Data Bank (PDB)^34^. We retrieved structures containing at least one RNA polymer entity and subsequently excluded those containing protein or DNA chains. Following the ligand composition criteria of HARIBOSS^29^, but without its molecular weight restriction, candidate ligands were defined as carbon-containing non-polymer components composed exclusively of C, H, N, O, P, S, F, Cl, Br, B, Si, Se and Co. Among the remaining structures, those with at least one ligand satisfying these criteria were classified as candidate holo structures. Candidate apo structures contained no qualifying ligand anywhere in the PDB file, although water, metal ions and other components that did not meet the ligand criteria were permitted. For each retained ligand, the binding pocket was defined as a set of RNA nucleotides containing at least one heavy atom within 8 Å of any ligand heavy atom, identified using a spatial neighbor search. The resulting RNA-ligand pockets were subsequently used as queries to identify sequence-matched ligand-free counterpart structures.

### Identification of apo-holo pairs

#### Sequence-based identification of counterpart chains

For each ligand-bound pocket, the full sequence of every RNA chain contributing to the binding site was reconstructed, with modified nucleotides mapped to their canonical parent bases and residues lacking a canonical parent represented as ‘N’. The same sequence-processing procedure was applied to all collected potential apo and holo structures, creating a searchable set of candidate structural counterparts. Pocket sequences were then queried against this database using BLASTN^35^ program. Candidate counterparts were retained only when BLASTN sequence identity over the aligned region, query coverage, and subject coverage were each ≥90%, thereby excluding poorly matched or partial-chain alignments. This BLASTN-based filtering criterion is distinct from the global sequence identity also reported in SMARTFlexDB webpage, which is calculated independently over the complete declared (SEQRES) RNA sequences using a Needleman-Wunsch global alignment^36^.

#### Pocket-region match and apo-holo assignment

Sequence-level filtering identified closely matched RNA chains but did not ensure conservation of the ligand-binding pocket. We therefore performed an additional pocket-level verification for each candidate match. The holo pocket was defined as the set of RNA nucleotides containing at least one heavy atom within 8 Å of a ligand heavy atom. Full-length query and candidate sequences were globally aligned, and each pocket nucleotide in the ligand-bound structure was mapped to the corresponding position in the candidate structure. A match was retained only when all pocket nucleotides could be mapped and were identical. During position-by-position comparison of pocket residues, positions containing ‘N’ in either sequence were compared using the original residue names rather than the one-letter codes. For single-chain pockets, a candidate apo structure was accepted if it contained one verified counterpart to the ligand-binding chain. For multi-chain pockets, a candidate apo structure was accepted only if it contained a distinct verified counterpart for every ligand-binding chain, ensuring that all chains contributing to the pocket were represented in the same structure. Structures containing the small-molecule ligand were included in SMARTFlexDB as the holo samples, whereas sequence- and pocket-matched structures were included as apo only when no qualifying ligand was present anywhere in the structure. For each unique apo-holo PDB pair, distinct binding pockets occupied by identical or different ligands in the holo RNA structure were presented within a single database entry, while retaining pocket-specific annotations and structural comparisons.

### RNA, ligand and interaction annotations

#### RNA identity and sequence

For each apo-holo comparison, SMARTFlexDB records the source PDB entries, the RNA chains contributing to the ligand-binding pocket, their normalized sequences, and an explicit nucleotide-level correspondence between the two states. Modified nucleotides were mapped to their canonical parent bases, with positions sharing the same parent base treated as sequence matches during apo-holo matching. Differences in deposited chemical component identities at these positions were annotated separately and highlighted in the sequence alignment on the web interface, with both component names displayed on hover. RNA functional types were assigned using entity-level annotations of the Nucleic Acid Knowledgebase (NAKB)^37^. Each RNA entity was assigned the most specific available functional class in the NAKB hierarchy. Structural annotations were not used for functional classification, and RNAs lacking a functional annotation were labelled as unannotated. For database browsing, fine-grained functional classes were grouped according to the first level of the NAKB functional hierarchy. The distribution of the 434 binding pockets across these broader functional categories is shown in Figure 1c, and the mapping between fine-grained classes and broader categories is provided in Supplementary Table S2.

#### Secondary structure and base-pair annotation

Secondary structures were annotated independently for the apo and holo states using DSSR^38^ program. For each structure, SMARTFlexDB stores the identified base pairs, dot-bracket representation and secondary-structure elements, including stems, hairpin loops, internal loops, bulges, junctions and single-stranded regions. Base pairs were annotated according to Leontis-Westhof geometry^39^ and were additionally classified as canonical or non-canonical based on DSSR pair assignments, with Watson-Crick and G-U wobble pairs considered canonical. Nucleotide-level mapping between the two states was then used to distinguish conserved base pairs from those detected only in the holo or apo structure.

#### Ligand identity and physicochemical properties

Ligands were identified by their PDB Chemical Component Dictionary (CCD) codes and deposited names. RDKit^40^ was used to calculate molecular weight, molecular formula, calculated logP, hydrogen-bond donor and acceptor counts, rotatable-bond count, aromatic-ring count, topological polar surface area and canonical SMILES. Each ligand card additionally provides a two-dimensional ligand depiction and links to external chemical databases (Supplementary Figure S1).

#### RNA-ligand atomic interaction and RNA pocket annotations

RNA-ligand interactions were annotated from the holo structures using the ligand-defined binding site described above. Hydrogen bonds were identified from atom-level donor-acceptor assignments generated by DSSR. π-stacking, halogen-bond and metal-coordination interactions were identified using PLIP^41^ in nucleic-acid receptor mode. Ligand burial was quantified from the holo structure using the Shrake-Rupley solvent-accessible surface-area algorithm^42^ implemented in Biopython^43^. Buried solvent-accessible surface area (SASA) was defined as the difference between the SASA of the isolated ligand and that of the same ligand in the context of the binding-site RNA. The latter calculation included the RNA chains contributing to the 8 Å binding site, while excluding water molecules, ions, free cofactors, additional ligand copies and RNA chains outside the binding-site context. The buried fraction was calculated by normalizing the buried SASA by the isolated-ligand SASA.

### Structural superposition and conformational analysis

#### Structural superposition

An explicit nucleotide-level correspondence was first established between the apo and holo structures by globally aligning their declared (SEQRES) RNA sequences using the Needleman-Wunsch algorithm. Only aligned positions containing identical nucleotides that were modelled in both structures were retained, and the apo RNA was then superposed onto the holo RNA by a least-squares Kabsch^44^ fit of the corresponding C3′ atoms. For multi-chain binding sites, matched nucleotides from the RNA chains contributing to the ligand-binding pocket were fitted simultaneously in a single superposition, after excluding chains identified as crystal-packing neighbors. Each holo chain was matched to at most one apo counterpart, and only admissible chain pairs were included in the fit, thereby preserving differences in the relative arrangement of the constituent chains. US-align was applied independently in RNA mode to assess overall fold similarity^45^. The resulting TM-score was normalized by the length of the holo ligand-host RNA chain. For visualization, a multi-frame morph was generated using custom code to linearly interpolate Cartesian coordinates between the superposed apo and holo structures, with Gemmi^46^ python package used for structural file input and output. The animation was rendered using Mol*^47^ and 3Dmol.js^48^. This geometric interpolation illustrates differences between the experimental endpoints and does not represent a molecular-dynamics trajectory or a physical ligand-binding pathway.

#### Global and local conformational descriptors

Global conformational change was quantified as the RMSD across all matched nucleotide pairs from the RNA chains included in the shared structural superposition. Overall fold similarity was independently described by the holo-length-normalized US-align TM-score. Local conformational change was quantified as the Euclidean displacement between each pair of matched apo and holo C3′ atoms. These values define the per-nucleotide displacement profile, which is reported from each apo-holo comparison. Pocket RMSD was calculated as the root-mean-square displacement of the matched holo-state pocket nucleotides across all RNA chains contributing to the binding site, measured in the shared structural-superposition frame. The holo pocket was defined as the set of RNA nucleotides containing at least one heavy atom within 8 Å of a ligand heavy atom. The radius of gyration ( *R*_g_ ) was calculated independently for the apo and holo structures using the matched nucleotide coordinates from all RNA chains included in the shared structural superposition, treating the combined coordinates as a single structural assembly. For *N* matched nucleotides, *R*_g_ was defined as *R*_g_ = 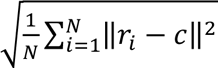, where *r_i_* is the C3′ coordinate of nucleotide *i* and 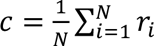 is the geometric centroid of the matched C3′ coordinates. All C3′ atoms were weighted equally. The change in radius of gyration was defined as Δ*R*_g_ = *R*_g_(ℎ*olo*) − *R*_g_(*apo*), such that negative values indicate compaction and positive values indicate expansion of the matched RNA structure upon ligand binding.

#### Base-pair rearrangements

Base pairs were annotated independently in the apo and holo structures using DSSR^38^ and mapped between the two states through the nucleotide-level correspondence. A base pair was classified as conserved when the same pair of mapped nucleotides was detected in both apo and holo structures, as formed when it was detected only in the holo structure, and as disrupted when it was detected only in the apo structure. SMARTFlexDB records the identities and total numbers of conserved, formed and disrupted base pairs, enabling ligand-associated secondary-structure rearrangements to be examined at nucleotide resolution.

### Clustering and multi-ligand pocket groups

Three complementary clustering schemes were used to support redundancy-aware exploration of SMARTFlexDB. The ligand-binding RNA chain sequences associated with each holo pocket were clustered with MMseqs2^49^ using minimum thresholds of 70% sequence identity and 80% coverage. The global folds of the corresponding host RNA chains were compared in an all-against-all manner with RMalign^50^. Structural clusters were defined as connected components of the graph formed by pairwise RMscore values ≥0.75. Pocket-level structural clustering followed the same procedure after restricting each RNA chain to nucleotides within 8 Å of the ligand. For visual exploration, the global-structure and pocket-structure distance matrices, defined as 1 - RMscore, were embedded in two dimensions using t-SNE^51^ , whereas the sequence projection was generated from a separately calculated k-mer distance matrix. In addition, holo-state RNA structures with closely matched RNA sequences and at least two distinct bound ligands were grouped into multi-ligand pocket groups (Supplementary Figure S7), enabling comparison of ligand-specific interactions and associated pocket conformational variations within a common RNA context.

### Database implementation

SMARTFlexDB was implemented using a Python-based FastAPI backend and a browser-based frontend developed in HTML, CSS and JavaScript. Interactive three-dimensional representations of the apo and holo structures, structural overlays and conformational morphs are rendered using Mol*^47^ and 3Dmol.js^48^. Per-chain RNA secondary structures are displayed using fornac^52^, while multi-chain joined views and base-pair rearrangement diagrams are rendered using custom SVG schematics. Two-dimensional ligand depictions are generated using RDKit. A REST API provides programmatic access to pair-level annotations, clustering and multi-ligand pocket groups information, coordinate files and other derived structural data. All retrieval endpoints are read-only, while a separate POST endpoint supports submission of user-defined apo-holo structure pairs for analysis. The query interface accepts user-submitted apo and holo structures in mmCIF format together with a ligand identifier. Submitted structures are validated and analyzed using a defined subset of the same annotation modules applied during database construction, and the resulting structural and conformational annotations are presented through the standard pair-page interface.

## RESULTS

### Current release and structural coverage

The current release contains 434 binding pockets from 266 holo structures. Each pocket is paired with all matching apo structures, yielding 1,187 pocket-level apo-holo comparisons organized into 734 apo-holo database entries. The collection comprises 452 distinct PDB structures, including 266 holo and 186 apo structures, and spans 150 ligands and 23 fine-grained RNA functional classes. At the broader functional level, riboswitches constitute the largest fraction of the dataset (166 of the 434 binding pockets, 38%), followed by RNAs involved in protein synthesis (110, 25%) and catalytic RNAs (84, 19%). Most binding pockets involve one RNA chain (276, 64%) or two RNA chains (153, 35%). The dataset also spans diverse secondary structure environments, with stems present in 423 binding pockets (97%), internal loops in 259 (60%) and multi-helix junctions in 219 (50%). In addition, 38 multi-ligand pocket groups capture distinct binding ligands within shared RNA contexts (Figure 1c). The data collection and curation workflow is summarized in Figure 1b, with detailed selection criteria and counts at each stage provided in Supplementary Table S1.

### SMARTFlexDB enables multidimensional apo-holo comparisons

Each SMARTFlexDB apo-holo pair page integrates sequence, three-dimensional structure, secondary structure, nucleotide-level conformational differences and RNA-ligand interactions within a unified interface (Figure 2; Supplementary Figure S1-S6). The sequence view combines a user-defined binding-site cutoff with apo-holo sequence alignment, nucleotide correspondence and the per-nucleotide displacement profile, enabling structural differences to be related directly to individual RNA positions (Figure 2a). The corresponding three-dimensional view superposes the apo and holo structures in a shared coordinate frame while retaining the bound ligand and allowing interactive control of molecular representation and binding-site highlighting (Figure 2b). A geometric apo-to-holo morph can be played interactively to visualize the structural difference between the two experimentally determined endpoints.

**Figure 2.**
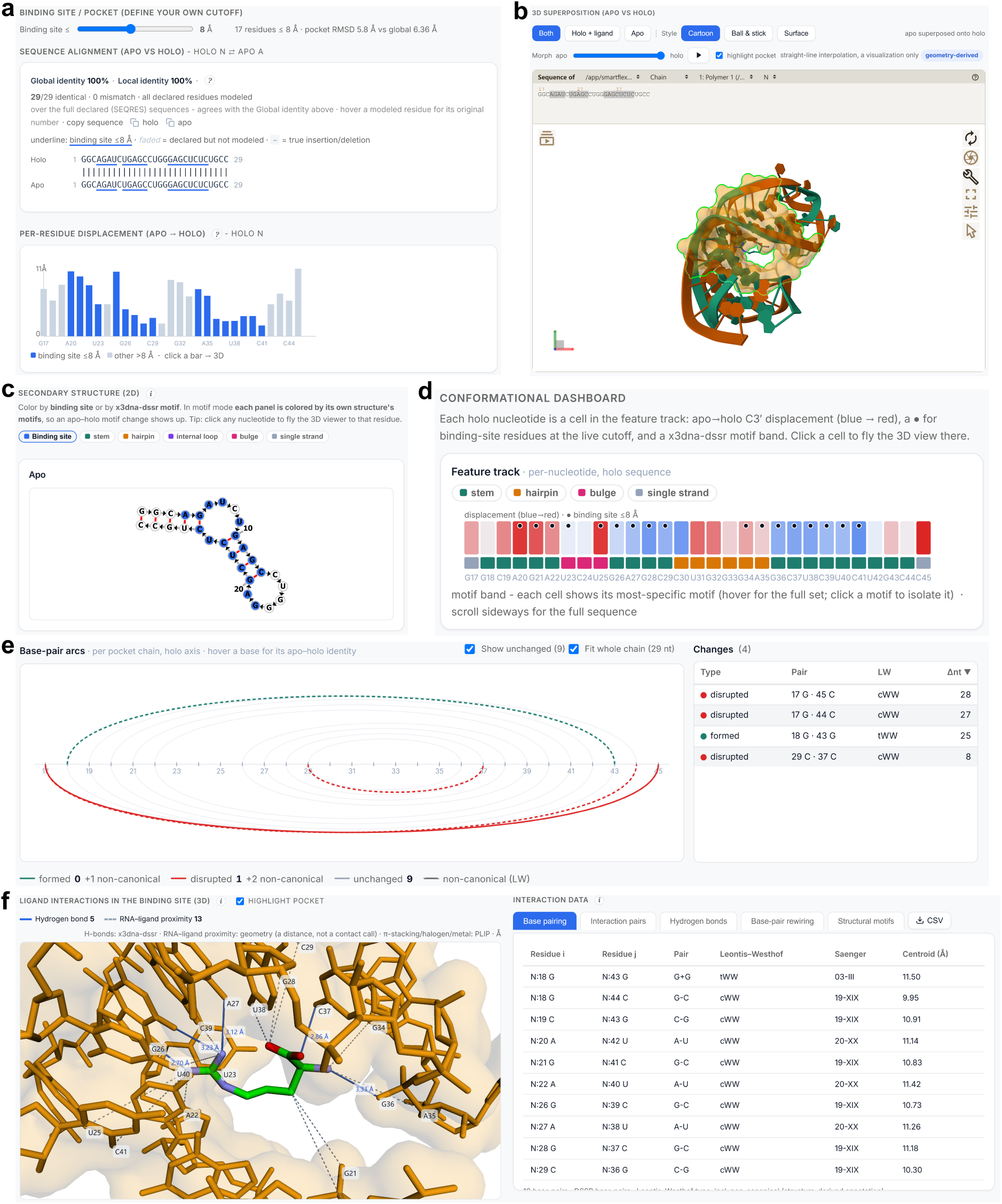
Multidimensional sequence, structure, and interaction analysis of a representative SMARTFlexDB apo-holo pair. **(a)** Binding-site definition, apo-holo sequence alignment and per-nucleotide displacement profile, with binding-site nucleotides highlighted along the aligned sequence. **(b)** Interactive three-dimensional superposition of the apo and holo RNA structures and the bound ligand. **(c)** Two-dimensional secondary-structure representation, colored by binding-site membership or by DSSR-derived structural motif. **(d)** Nucleotide-level conformational dashboard aligning apo-holo displacement, binding-site membership and DSSR-derived structural motifs along the holo RNA sequence. **(e)** Base-pair rearrangement analysis showing conserved, formed and disrupted base pairs between the apo and holo states, with nucleotide-level pair identities and Leontis-Westhof annotations. **(f)** Ligand binding-site interaction view combining three-dimensional visualization of RNA-ligand contacts with an interactive annotation interface for binding-site interactions, hydrogen bonds, base pairing, base-pair rearrangements and structural motifs.

Secondary structure annotations provide a complementary two-dimensional representation of the RNA and identify the structural motifs associated with individual nucleotides (Figure 2c). These annotations are integrated with a nucleotide-level conformational dashboard that aligns structural-motif assignment, binding-site membership and apo-holo displacement along the holo RNA sequence (Figure 2d). Base-pair rearrangements are displayed separately as conserved, formed and disrupted interactions between mapped nucleotides, together with nucleotide identities and Leontis-Westhof annotations (Figure 2e). These linked views allow global and local structural differences to be traced from the three-dimensional comparison to specific nucleotides, motifs and base-pair changes.

The ligand binding-site view places the bound ligand within its RNA environment and visualizes annotated RNA-ligand contacts in three dimensions (Figure 2f). The accompanying annotation interface provides complementary nucleotide-level information on binding-site contacts, hydrogen bonds, base pairing, base-pair rearrangements and structural motifs. Together, these representations connect the interactions observed in the deposited holo structure with structural differences between the experimentally determined apo and holo endpoints.

### Clustering and multi-ligand pocket groups

SMARTFlexDB provides cluster assignments based on RNA sequence, global structure and pocket structure (Supplementary Figure S9). Under the thresholds described above, the current release contains 95 sequence clusters, 100 structure clusters and 102 binding-pocket clusters. Interactive t-SNE projections allow users to locate related entries and export cluster assignments. These complementary labels support redundancy-aware analysis at the sequence, fold and binding-pocket levels.

The multi-ligand pocket group view displays ligands observed with the same or closely related RNA pocket context, enabling direct comparison of ligand-specific recognition patterns (Figure 3a; Supplementary Figure S7 and S8). The current release contains 38 multi-ligand pocket groups, with the largest group containing 23 distinct ligands. Each group summarizes ligand structures and physicochemical properties, pocket RMSD and the RNA nucleotides contacted by each ligand, thereby enabling direct comparison of ligand recognition within a shared RNA context.

**Figure 3.**
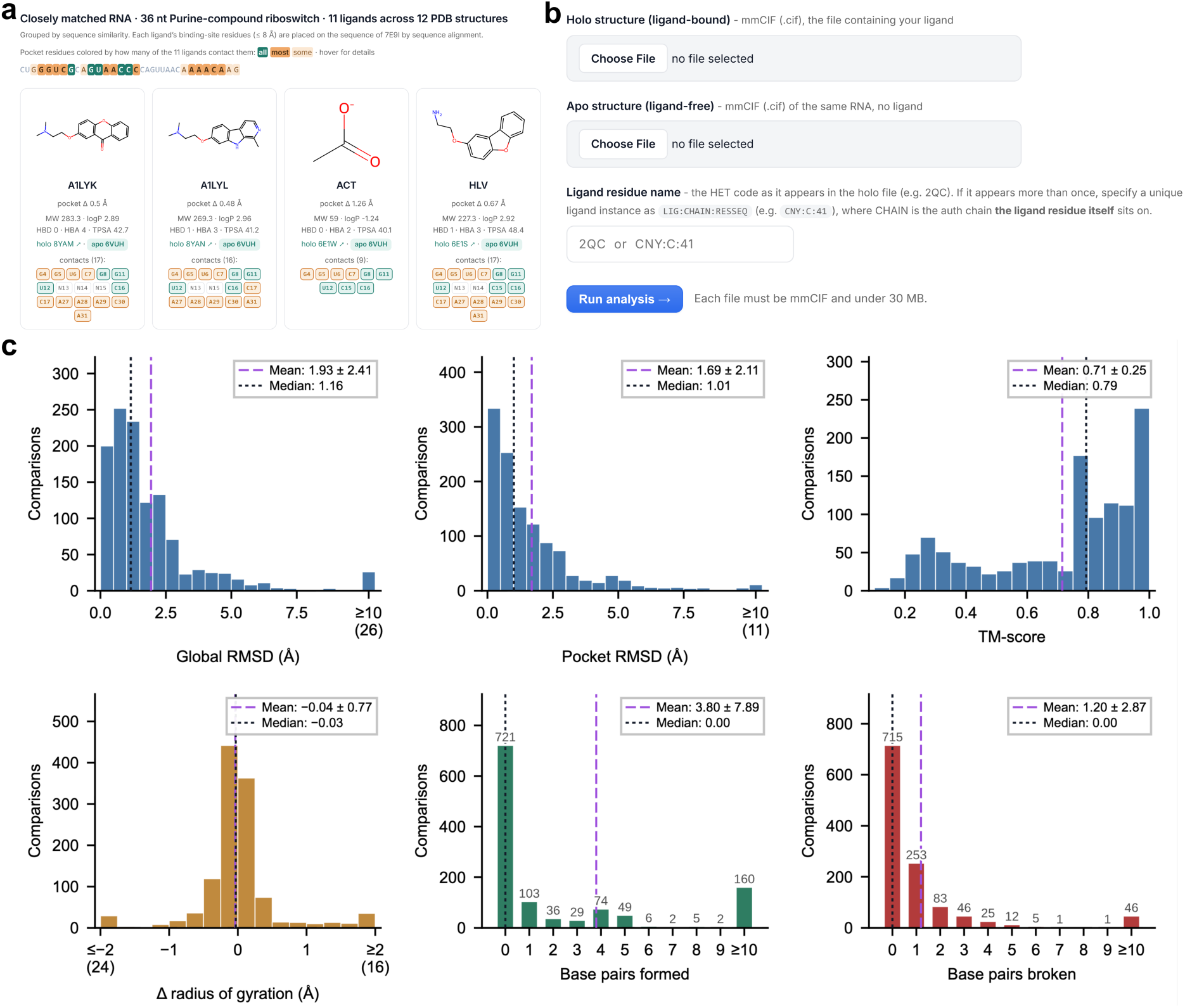
SMARTFlexDB dataset overview and analysis interfaces. **(a)** Representative multi-ligand pocket group, showing the ligands observed with the same or closely related RNA scaffold. Ligand structures and physicochemical properties are shown alongside pocket RMSD and contacted RNA nucleotides, enabling comparison of ligands within a shared RNA context. **(b)** Query interface for analysis of user-supplied apo and holo RNA structures. Users upload the two structures in mmCIF format and specify the ligand identifier in the holo structure; the submitted pair is then processed through the supported structural-comparison and annotation workflow. **(c)** Distributions of global RMSD, pocket RMSD, TM-score, change in radius of gyration (ΔRg), and the numbers of canonical base pairs formed or disrupted between the apo and holo states across the 1,187 pocket-level apo-holo comparisons of the release. Purple and black dashed lines denote the mean and median, respectively; legends report the mean ± s.d. and median. ΔRg is defined as Rg(holo) − Rg(apo).

### Query service, downloading and API

The query page extends the SMARTFlexDB analysis workflow to user-supplied structures. Users can upload apo and holo structures in mmCIF format and specify the ligand identifier in the holo structure (Figure 3b). Following input validation, the server establishes nucleotide-level sequence correspondence, performs structural superposition and calculates conformational descriptors, secondary-structure changes and RNA-ligand interactions. For each submitted pair, the ligand-binding holo RNA chain is also searched against holo RNA sequences in the current SMARTFlexDB release using MMseqs2, and the query is assigned the cluster label of the best-matching sequence satisfying ≥70% sequence identity and ≥80% sequence coverage, when such a match is available. Results are presented through the same pair-page interface used for curated records.

All release data are accessible without registration. The bulk table (*smartflexdb_pairs.tsv*, available from the API page) contains 1,187 rows, one per pocket-level apo-holo comparison (Supplementary Note S2). These group into the 734 apo-holo database entries shown on the website. The API provides access to curated binding-pocket records listing every apo structure each pocket is compared with, classification and multi-ligand pocket group information, structural files and downloadable record-level bundles. API documentation provides reusable examples for programmatic access, and record-level archives contain the corresponding coordinate files and derived annotations for independent downstream analysis.

### Conformational characteristics

Across the 1,187 pocket-level apo-holo comparisons, SMARTFlexDB captures a broad range of apo-holo structural differences (Figure 3c). Global RMSD had a mean of 1.93 ± 2.41 Å and a median of 1.16 Å, while pocket RMSD was slightly smaller, with a mean of 1.69 ± 2.11 Å and a median of 1.01 Å. Thus, structural differences at the ligand-binding site are, at the dataset level, comparable in magnitude to those observed across the overall RNA structure. TM-score provided a complementary measure of fold similarity, with a mean of 0.71 ± 0.25 and a median of 0.79.

Changes in radius of gyration were centred near zero (mean ΔRg = -0.04 ± 0.77 Å; median = -0.03 Å), indicating no strong overall tendency towards either compaction or expansion between the deposited apo and holo states. Canonical base-pair rearrangements were comparatively sparse: 721 of 1,187 comparisons formed no new canonical base pairs and 715 broke none, with median values of zero for both measures. Together, these distributions show that SMARTFlexDB spans substantial three-dimensional endpoint variability while extensive canonical base-pair rewiring is not a universal feature of apo-holo structural differences.

## DISCUSSION

RNA-targeted small molecules offer a promising therapeutic strategy for diseases that remain inaccessible to protein-targeting approaches. RNA-focused libraries and databases are therefore essential for advancing RNA-targeted drug discovery. However, existing resources either lack three-dimensional structural information (SM2miR^24^, Inforna 2.0^25^, NoncoRNA^26^, R-BIND 2.0^27^), or are limited to static RNA-ligand structural snapshots, as in HARIBOSS and R-SIM^28^, despite the central role of RNA conformational plasticity in ligand recognition. SMARTFlexDB addresses this gap by linking apo and holo RNA structures at both the global and binding-pocket levels and quantitatively characterizing their structural differences, RNA properties, and ligand features. By making the apo-holo comparison the primary, consistently annotated record, SMARTFlexDB enables RNA-small molecule recognition to be analyzed beyond conventional individual complex structures through direct comparisons between ligand-free and ligand-bound states. Through an interactive web interface, users can examine detailed conformational changes, compare secondary structures and RNA motifs, and upload self-paired apo-holo structures for customized analysis.

SMARTFlexDB is designed to support multiple research communities. For RNA and structural biologists, it provides an experimentally grounded resource of ligand-associated RNA conformational changes, including sequence alignments, secondary structure comparisons, RNA motif annotations, three-dimensional overlays, and pocket-level remodeling metrics. These features enable users to examine how nucleotide displacement, base-pairing changes, and local pocket rearrangements contribute to RNA structural flexibility and molecular recognition. SMARTFlexDB quantifies structural differences between experimentally determined apo and holo RNA structures; however, these differences should not necessarily be interpreted as being caused by ligand binding. Intrinsic RNA conformational heterogeneity, differences in nucleotide modification state, and experimental context may also contribute to apo-holo variation, and mechanistic interpretation therefore remains structure- and context-dependent. Where multiple matched structures are available, comparison across alternative apo or holo conformations can provide additional context for assessing the consistency of observed structural changes.

For medicinal and computational chemists, SMARTFlexDB provides pocket- and ligand-level annotations directly relevant to RNA-targeted small molecule discovery. RNA pocket descriptors, RNA-ligand atomic interactions, and quantitative measures of pocket flexibility can inform druggability assessment, flexible and ensemble docking, and structure-guided optimization. Because RNA targets are often available only in ligand-free conformations during early discovery, paired apo-holo comparisons provide important insights into cryptic pocket identification and how small molecules exploit RNA structural plasticity. Multi-ligand pocket groups, which collect different ligands bound to a same or a closely related RNA pocket further enable identification of key interactions and comparison of ligand-specific recognition patterns and pocket changes.

SMARTFlexDB also provides a standardized foundation for computational method development and evaluation. As interest grows in RNA-focused machine-learning and physics-based approaches for RNA-ligand interaction prediction, docking, virtual screening, and affinity estimation, most existing methods are developed and evaluated primarily on the fixed holo RNA conformations. However, RNA conformational flexibility and binding-associated structural changes can substantially limit their performance in realistic drug-discovery settings. The experimentally grounded, matched apo-holo pairs in SMARTFlexDB enable flexible docking evaluation, binding site and druggability prediction from ligand-free structures, assessment of pocket-aware generative models under conformational flexibility. Sequence-, structure-, and pocket-level clustering further supports leakage-controlled data splits for fair and reproducible benchmarking.

With rapid advances in RNA-targeted ligand discovery and high-resolution RNA and RNA-ligand structure determination, we will continue to update SMARTFlexDB regularly by expanding its data coverage, enriching annotations, and improving the functionality of the web interface. Future extensions of SMARTFlexDB may incorporate additional structural and biophysical annotations, including RNA backbone conformational descriptors^53^, rigid-base coordinate frames for quantitative analysis of base orientation and stacking, and experimentally supported binding thermodynamic information where data are available. These additions could provide a more detailed characterization of RNA structural changes and further connect conformational differences with molecular recognition and binding energetics. For holo RNA structures with multiple binding pockets, SMARTFlexDB enables joint analysis of these sites within the same apo-holo context and may support future investigation of inter-pocket coupling and potential allosteric relationships. Future releases will incorporate automated synchronization with new PDB depositions with necessary manual quality control. Although the current release focuses on experimentally determined RNA-small molecule complexes, future versions may incorporate computationally generated RNA or RNA-ligand structures with appropriate confidence or fidelity labels as RNA structure prediction and co-folding models continue to improve. We also plan to extend SMARTFlexDB to RNA complexes containing protein or DNA components, thereby broadening its coverage of biologically relevant RNA-containing assemblies. We expect SMARTFlexDB, as a dedicated and systematically organized database of apo-holo RNA structures, to provide substantial value to the rapidly growing field of structure-based RNA-targeted drug discovery.

## ACKNOWLEDGEMENTS

We thank Zekun Yang (University of Florida) for initial data collection and technical discussions. The graphical abstract was created in BioRender. Jiang, S. (https://BioRender.com/hr8a6nu) is licensed under CC BY 4.0.

## Author contributions

All authors contributed to writing - review & editing. Y.L.: conceptualization, investigation, methodology, supervision, writing – original draft. S.J., J.W., and A.T.: methodology, data curation, formal analysis, investigation, writing – original draft. S.J., J.W.: software. G.S., J.L.C., C.L., and M.D.D.: investigation, writing – original draft. All authors reviewed and approved the final version.

## DATA AVAILABILITY

SMARTFlexDB is freely accessible without registration at https://aidd.rc.ufl.edu/app/smartflexdb/. API documentation and programmatic access are provided at https://aidd.rc.ufl.edu/app/smartflexdb/api.html. Pair-level annotation bundles and coordinate files can be downloaded from the corresponding entry pages. The curated experimental structures are available from the RCSB Protein Data Bank under the accession codes recorded in SMARTFlexDB.

